# The glyoxylate shunt protein ICL-1 protects from mitochondrial superoxide stress through activation of the mitochondrial unfolded protein response

**DOI:** 10.1101/2022.10.10.511563

**Authors:** Guoqiang Wang, Ricardo Laranjeiro, Stephanie LeValley, Jeremy M. Van Raamsdonk, Monica Driscoll

## Abstract

Eliminating mitochondrial superoxide dismutase (SOD) causes neonatal lethality in mice and death of flies within 24 hours after eclosion. Deletion of mitochondrial *sod* genes in *C. elegans* impairs fertility as well, but surprisingly is not detrimental to survival of progeny generated. The comparison of metabolic pathways among mouse, flies and nematodes reveals that mice and flies lack the glyoxylate shunt. Here we show that ICL-1, the only protein of the glyoxylate shunt, is critical for protection against embryonic lethality resulting from elevated levels of mitochondrial superoxide. In exploring the mechanism by which ICL-1 protects against ROS-mediated embryonic lethality, we find that ICL-1 is required for the efficient activation of mitochondrial unfolded protein response (UPR^mt^) and that the UPR^mt^ is essential to suppress embryonic/neonatal lethality in animals lacking mitochondrial SOD. In sum, we identified a biochemical pathway that highlights a molecular strategy for combating superoxide consequences in cells.

## INTRODUCTION

Continuous balance of energy conversion and consumption is key for living organisms, with efficient mitochondrial energy processing absolutely required for animals with high energy demands. The mitochondrial electron transport chain (ETC) plays a critical role in energy processing, but electrons can occasionally escape the ETC to be captured by oxygen to form superoxide, a highly reactive negatively charged oxidant that can damage cell components. To protect against the superoxide threat to mitochondrial function, superoxide dismutase (SOD), especially mitochondrial MnSOD, serves as a critical defense for viability (Duttaroy et al., 2003; Lebovitz et al., 1996; Li et al., 1995).

Unlike other model systems like mouse and fly in which mitochondrial SOD is critical for development and survival (Duttaroy *et al*., 2003; Lebovitz *et al*., 1996; Li *et al*., 1995), *C. elegans* can overcome the loss of mitochondrial SOD to develop to adult, and mild superoxide stress that accompanies mitochondrial *sod-2* deletion can even extend lifelong survival (Van Raamsdonk and Hekimi, 2009). The distinctive contrast in the developmental consequences of mitochondrial SOD disruption between *C. elegans* and other organisms suggests the possibility that *C. elegans* features a protective pathway counteracting mitochondrial superoxide stress that is lacking in fly and mouse.

The conservation of many mitochondrial genes and pathways with mammalian mitochondria and the numerous genetic tools applicable to experimentation make *C. elegans* a powerful model organism in which to study mitochondrial biology (van der Bliek et al., 2017). RNAi knockdowns or loss of function mutations in genes encoding key components of the ETC are often associated with both a surge of oxidative stress and activation of the glyoxylate shunt in *C. elegans* (Bennett et al., 2017; Gallo et al., 2011; Hekimi et al., 2016; Morgan et al., 2015; Pujol et al., 2013; Senchuk et al., 2018; Zuryn et al., 2010), suggesting a potential regulatory relationship between oxidative stress and the glyoxylate shunt.

The glyoxylate shunt requires two enzymatic activities: isocitrate lyase and malate synthase. Both activities are mediated by the *C. elegans* bifunctional enzyme ICL-1. As a branch of the tricarboxylic acid (TCA) cycle, the glyoxylate shunt consumes an acetyl-CoA while converting isocitrate directly to malate and succinate (Figure 1A). The glyoxylate shunt results in the bypass of oxidative decarboxylation steps of the canonical TCA cycle (Dolan and Welch, 2018), such that two steps of regulated NADH production are skipped and 2 NADH produced by the full TCA cycle are not generated). Genomic evidence suggests that the glyoxylate shunt is present in bacteria, protists, plants, fungi and nematodes (Kondrashov et al., 2006). Mammals appear to have lost the glyoxylate shunt during evolution. Strikingly, however, transgenic restoration of the glyoxylate shunt into mouse liver by adding back isocitrate lyase and malate synthase can confer resistance to diet-induced obesity (Dean et al., 2009), revealing just one potential therapeutic application of elaborating how the glyoxylate shunt engages basic cell metabolism.

**Figure 1.**
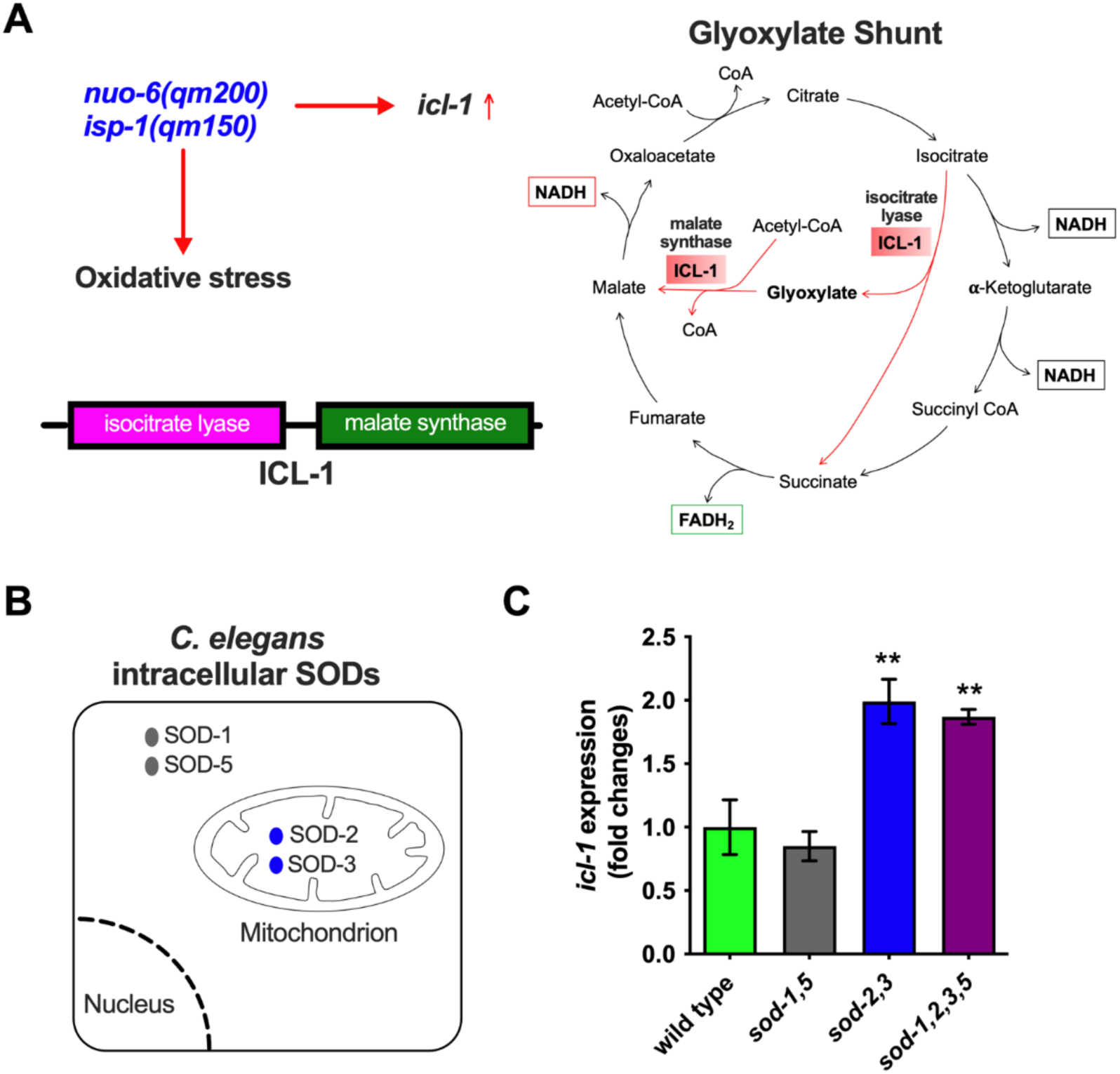
The induction of *icl-1* gene expression is specific to mitochondrial superoxide stress. A. Specific ETC (electron transport chain) point mutants, *nuo-6(qm200)* or *isp-1(qm150)*, can induce *icl-1* gene expression (Senchuk *et al*., 2018) and are correlated with oxidative stress (van der Bliek *et al*., 2017). ICL-1 is a bifunctional enzyme with both isocitrate lyase activity and malate synthase activity. A simplified diagram of the tricarboxylic acid (TCA) cycle with a bypass through the glyoxylate shunt highlighted. B. A simplified diagram summarizing the subcellular location of SOD proteins in *C. elegans* cells. Both SOD-1 and SOD-5 are located in cytosol, while SOD-2 and SOD-3 are located in the mitochondrial matrix (Van Raamsdonk and Hekimi, 2012). C. RT-qPCR quantitation of *icl-1* gene expression at the L4 stage of wild type, the *sod-1 sod-5* double null mutant, the *sod-2 sod-3* double null mutant or the *sod-1 sod-2 sod-3 sod-5* quadruple null mutant (n = 3 independent trials, ** *p*<0.01 as compared to wild type in one-way ANOVA).

In *C. elegans*, knocking out the primary mitochondrial SOD gene, *sod-2*, increases *C. elegans* lifespan (Van Raamsdonk and Hekimi, 2009) and activates the mitochondrial unfolded protein response (UPR^mt^) (Wu et al., 2018), a conserved transcriptional response to mitochondrial dysfunction (Shpilka and Haynes, 2018). ATFS-1, the key transcriptional factor mediating the UPR^mt^ (Nargund et al., 2012), can directly bind to the promoter region of *icl-1*, as shown by the ATFS-1 ChIP-seq data (Nargund et al., 2015). A constitutively active (CA) mutation in *atfs-1* significantly increases the expression of *icl-1* (Lin et al., 2016). Taken together, these observations suggest a tight relationship between mitochondrial superoxide stress, ATFS-1-mediated UPR^mt^ activation and the induction of the glyoxylate shunt to protect against oxidative stress.

Here we report that *icl-1* expression is specifically induced by mitochondrial superoxide stress but not cytosolic superoxide stress. The induction of *icl-1* protects against mitochondrial superoxide stress allowing for embryonic survival under conditions of elevated mitochondrial superoxide. We also find that full activation of the UPR^mt^ is dependent on ICL-1 and that constitutive activation of UPR^mt^ can bypass the requirement of ICL-1 for embryonic survival in the absence of mitochondrial SODs. Overall, our study provides new insight into how an ancient metabolic pathway plays an invaluable role in counteracting mitochondrial superoxide stress.

## RESULTS

### Eliminating mitochondrial SODs but not cytosolic SODs elevates *icl-1* transcript levels

Previous studies have shown that mutations affecting the mitochondrial ETC, such as *nuo-6(qm200)* or *isp-1(qm150)*, can induce *icl-1* expression (Senchuk *et al*., 2018) (Fig. 1A). As these mutations also increase superoxide production from complex I or III, respectively (van der Bliek *et al*., 2017), we wondered whether elevation of mitochondrial superoxide is sufficient in upregulate *icl-1*.

The *C. elegans* genome encodes five *sod* genes: *sod-1* and *sod-5* encode the cytosolic SODs; *sod-2* and *sod-3* encode the mitochondrial SODs; and *sod-4* encodes extracellular SOD (Fig. 1B). To determine if elevation of mitochondrial superoxide levels would increase the levels of *icl-1*, we quantitated *icl-1* expression in double null mutants lacking all mitochondrial SOD or all cytosolic SOD. Using quantitative reverse transcription PCR (RT-qPCR), we found that elimination of mitochondrial SODs significantly increases *icl-1* gene expression, while eliminating cytosolic SODs has no impact on *icl-1* gene expression. Eliminating both cytosolic and mitochondrial SODs has an impact on *icl-1* induction similar to eliminating only mitochondrial SODs (Fig. 1E). Together, these data indicate that the induction of *icl-1* gene expression is specific to mitochondrial superoxide stress.

### *icl-1* protects embryonic development when mitochondrial SODs are deleted

Although mutants lacking mitochondrial SODs exhibit normal (*sod-3*) or even extended lifespans (*sod-2*) (Van Raamsdonk and Hekimi, 2009), eliminating mitochondrial SODs causes about 70% reduction in brood size (Van Raamsdonk and Hekimi, 2009). Because the glyoxylate shunt should lower the superoxide levels (a deleterious factor to healthy reproduction) by lowering the generation of NADH, and thus might protect from superoxide toxicity by lowering NADH levels, we tested the impact of *icl-1(D)* on brood size.

We first used a simple plate assay that can reveal basic *C. elegans* progeny production levels. Rearing a single L4 hermaphrodite on the OP50 bacteria seeded plate for 7 days normally results in two generations that populate the culture plate (Fig. 2A, wild type). We find that the *icl-1* deletion mutant and the *sod-2; sod-3* double null mutant populate the plates well in this general assay (Fig. 2A), although as expected from previous work the *sod-2; sod-3* double null mutant, development is modestly delayed. In contrast, the *sod-2; icl-1; sod-3* triple null mutant produces only a very few progeny by day 7 (Fig. 2A), suggesting that in the absence of mitochondrial *sod, icl-1* is needed to enable progeny production.

**Figure 2.**
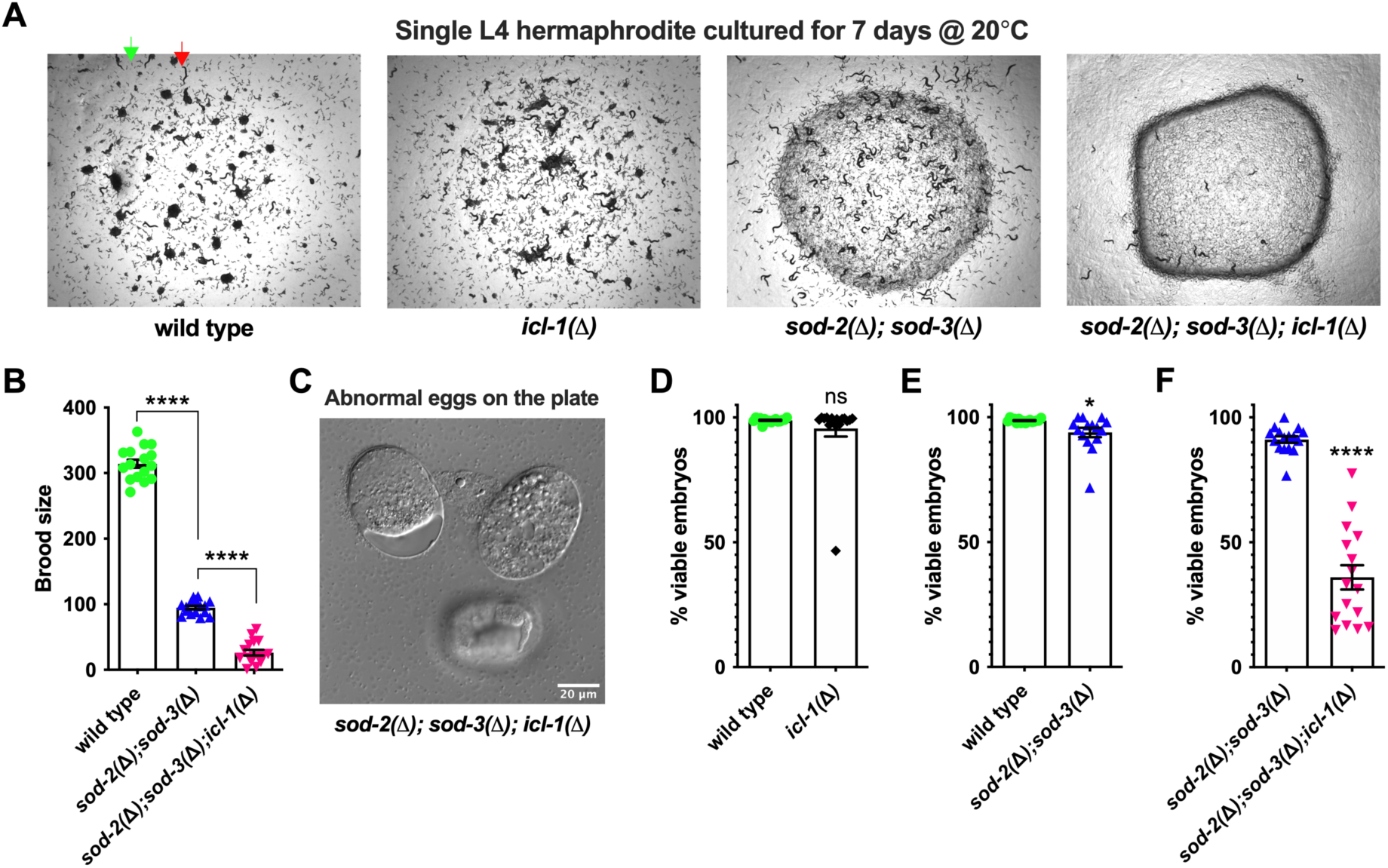
*icl-1* is required for the embryonic viability of mitochondrial *sod* mutant. A. Representative pictures of the outcome for 7-day culture of a single L4 hermaphrodite in a plate for strains indicated. The dark dots (green arrow) and the dark short lines (red arrow) are all progeny from a single L4 hermaphrodite; the circular-shape area is where the bacteria OP50 were seeded on the plate (light appearance for wild type or *icl-1* mutant; dark for others). B. Brood size (viable progeny of a single worm in its whole life) of the *sod-2 sod-3* double null mutant and *sod-2; icl-1; sod-3* triple null mutant (n = 16 biological repeats, **** *p*<0.0001 in oneway ANOVA). C. Representative DIC pictures showing dead/abnormal embryos of eggs collected from the plate of the *sod-2; icl-1; sod-3* triple null mutant. D. Percentage of viable embryos from wild type and *icl-1* null mutant (n = 16 biological repeats, *p*=0.3318 in unpaired two-tail *t-*test). E. Percentage of viable embryos from wild type and *sod-2; sod-3* double null mutant (n = 15 or 16 biological repeats, * *p*=0.0126 in unpaired two-tail *t-*test). F. Percentage of viable embryos from the *sod-2; sod-3* double null mutant and *sod-2; icl-1; sod-3* triple null mutant (n = 16 biological repeats, **** *p*<0.0001 in unpaired two-tail *t-*test).

We then quantitatively compared progeny production in the *sod-2; sod-3* double null mutant vs. the *sod-2; icl-1; sod-3* triple null mutant. We find that the triple *sod-2; icl-1; sod-3* triple mutant has a significantly reduced brood size as compared to the *sod-2 sod-3* double null mutant (Fig. 2B). We conclude that *icl-1* exerts a protective function for progeny production in compromised *sod-2; sod-3* double null mutants.

During the process of crossing the *icl-1* null mutant, with the *sod-2; sod-3* double null mutant, we uncovered deficits in recovery of the triple homozygous null mutant from *sod-2(-/-); icl-1(+/-); sod-3(-/-)*, which normally would be expected to be 25% of cross progeny, but only was recovered in ~7% of progeny. Low proportions of the triple null mutant suggest potential partial synthetic lethality. Indeed, examination of eggs produced by the triple null mutant revealed many embryos arrested at various stages of development that failed to hatch (examples in Fig. 2C).

To confirm developmental consequences of *icl-1* elimination on *sod-2; sod-3* embryonic viability, we quantitated hatching and survival. Although the *icl-1(D)* mutant exhibits a marginal reduction in brood size on its own (Fig. 1s), we found that *icl-1(D)* exhibits embryonic viability (percentage of hatched eggs among all eggs in the life of a single hermaphrodite) similar to wild type (Fig. 2D, 96% *icl-1(D)* vs. 99% WT). Likewise, the *sod-2; sod-3* double null mutant exhibits mildly reduced embryonic viability as compared to the wild type (Fig. 2E, 94% vs. 99%, *p*=0.0126). In contrast, the *sod-2; icl-1; sod-3* triple null mutant exhibits significantly diminished embryonic viability as compared to the *sod-2; sod-3* double null mutant (Fig. 2F, 36% triple mutant vs. 91% *sod-2; sod-3*).

We conclude that *icl-1*, and likely the glyoxylate shunt that ICL-1 catalyzes, is required for successful embryonic development when mitochondrial SODs are absent and superoxide is high.

### Elimination of ICL-1 does not markedly change mitochondrial membrane potential or paraquat sensitivity in the *sod-2 sod-3* background

The glyoxylate shunt shortcut of the full TCA cycle (Fig. 1A) could lead to extensive remodeling of mitochondrial metabolism and function. To assess general mitochondrial health when mitochondrial *sods* and/or *icl-1* are absent, we measured overall mitochondrial membrane potential (Δ**Ψ**_m_) in *sod; icl-1* and triple mutants. We stained hermaphrodites with the mitochondrial membrane potential dye TMREE, by feeding animals bacteria mixed with 100 μM TMREE for 24 hours. We took images for analysis immediately after removing the animals (day 1 adult) from the dye. We find that the *icl-1* null mutation only causes a trend of slight reduction in either whole-body (Fig. 3A) or oocyte Δ**Ψ**_m_ (Fig. 3B). In contrast, the mitochondrial *sod* double null mutations significantly reduce mitochondrial membrane potential in both whole-body measurement (Fig. 3A) and in oocyte measurement (Fig. 3B) as compared to the wild type. We find that the triple *sod-2; icl-1; sod-3* mutant is not significantly different from the *sod-2; sod-* 3 double mutant for whole-body Δ**Ψ**_m_ (Fig. 3A), and note a trend toward slight reduction in oocyte Δ**Ψ**_m_ (Fig. 3B, not statistically significant). Thus, *icl-1* deficiency does not further increase the mitochondrial membrane potential deficits associated with elimination of all mitochondrial SOD levels. The brood size boost provided by *icl-1* is thus likely to operate outside of adjusting the mitochondrial membrane potential deficits.

**Figure 3.**
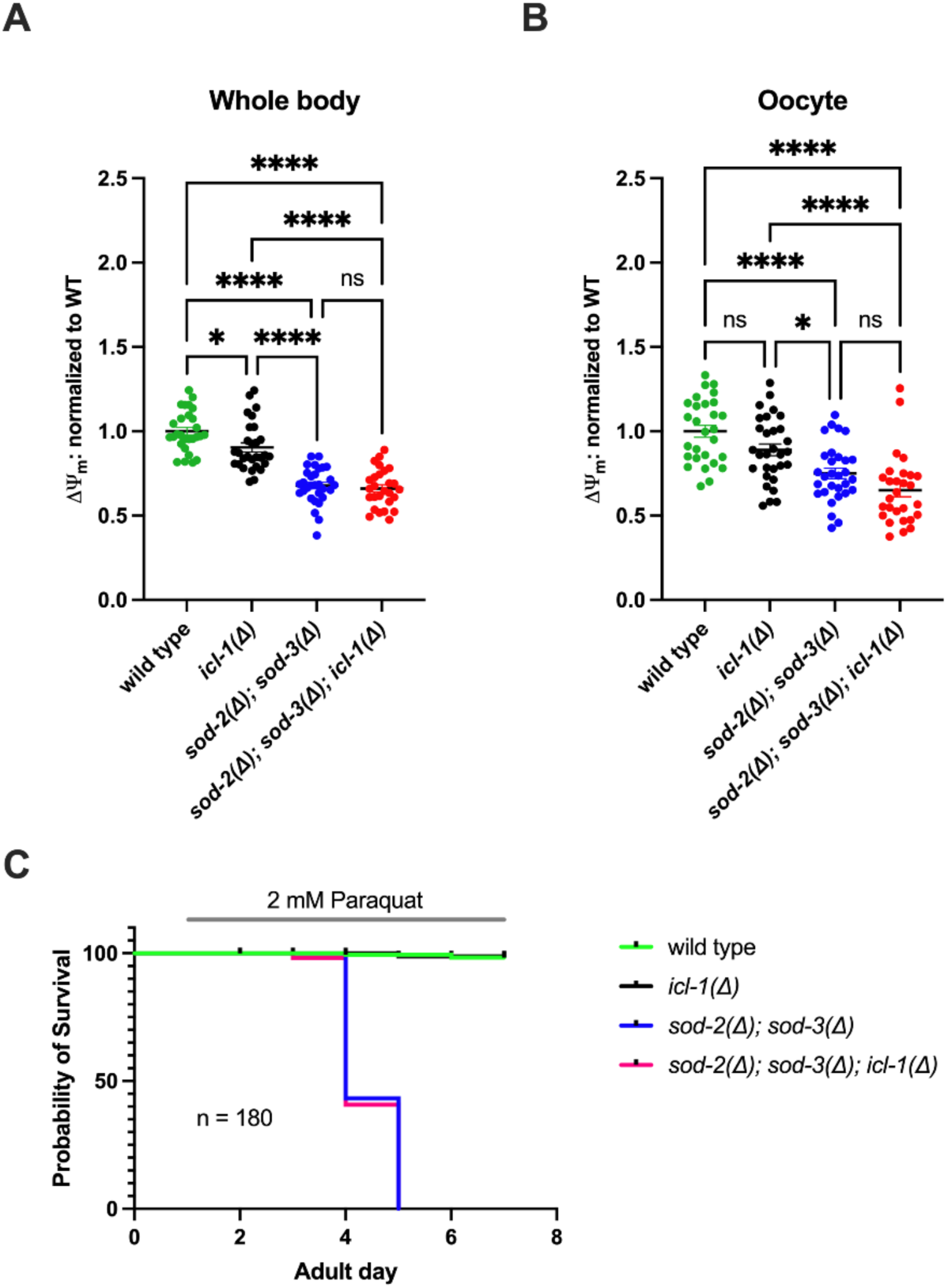
Mitochondrial membrane potential and paraquat sensitivity of the *sod-2; sod-3* mutant are not changed by eliminating ICL-1. A. Measurement of the TMREE (Tetramethylrhodamine, Ethyl Ester) fluorescence intensity (of the intact whole animal) to determine the relative mitochondrial membrane potential (Δ**Ψ**_m_) of animals that had been fed with 100 μM TMREE for 24 hours (from L4 stage) as normalized to the wild type. Total of ~ 45 hermaphrodites for each genotype from three independent trials. **** *p* < 0.0001; * *p* < 0.05 or ns (not significant) in one-way ANOVA. B. Measurement of the TMREE fluorescence intensity of oocytes in the gonad to determine the relative mitochondrial membrane potential (Δ**Ψ**_m_) of oocytes in animals that had been fed with 100 μM for 24 hours (from L4 stage) and normalized to the wild type. Total of ~45 hermaphrodites from three independent trials. **** *p* < 0.0001; * *p* < 0.05 or ns (not significant) in one-way ANOVA. C. Survival in the presence of paraquat (2 mM) in age-synchronized hermaphrodites. The treatments started from adult day 1 and the life status of each animal was recorded every 24 hours. Total of 180 hermaphrodites from 3 independent trials for each genotype; *p* < 0.0001 (wild type vs. *sod-2(Δ); sod-3(*Δ)) in Log-rank (Mantel-Cox) test.

To further test models of how ICL-1 might counteract mitochondrial superoxide stress, we quantitated mutant resistance to paraquat, a toxin that can increase mitochondrial superoxide production (Yamada and Fukushima, 1993). Wild type and *icl-1* null mutants are similarly resistant to paraquat toxicity at the administered 2mM paraquat dose (Fig. 3C), an indication that *icl-1* is not normally critical for superoxide stress resistance under paraquat exposure. The mitochondrial *sod* null mutants survive until adult day 4,5, when viability drops to near zero. Elimination of the *icl-1* gene in the *sod-2; sod-*3 mutant background neither accelerates nor delays the paraquat-induced *sod-2; sod-3* mutant death (Fig. 3C), suggesting that protective effects of *icl-1* deletion are unlikely to be anchored in superoxide detoxification.

Overall, ICL-1 appears dispensable for mitochondrial function and superoxide detoxification at least early in life, however, negative results of the assays described could reflect limited sensitivity of measurement. To probe further, we tested conditions that genetically perturb mitochondrial function and health. The mitochondrial unfolded protein response (UPR^mt^) acts as a native sensor of mitochondrial stress, including superoxide stress (Wu *et al*., 2018). We therefore tested UPR^mt^ activity as a readout to evaluate the impact of ICL-1 on mitochondrial superoxide stress.

### ICL-1 is required for an efficient activation of UPR^mt^

Eliminating mitochondrial SODs can lead to severe damage to mitochondrial function (Melov et al., 1999; Van Raamsdonk and Hekimi, 2009) and activation of the UPR^mt^ (Wu *et al*., 2018). The UPR^mt^ upregulates *icl-1* (Lin *et al*., 2016; Nargund *et al*., 2015), which our data suggest is critical for functional mitochondrial superoxide stress protection in development.

To better understand the relationship between ICL-1, the UPR^mt^, and toxicity associated with *sod-2; sod-3* progeny, we used the *hsp-6p::GFP* UPR^mt^ reporter strain to quantify UPR^mt^ activation in *sod-2;sod-3* mutants with and without ICL-1. We hypothesized that eliminating *icl-1* would push the *sod-2; sod-3* mutant to a higher mitochondrial stress and activate a stronger UPR^mt^ (Fig. 4A). However, we found that elimination of *icl-1* reduces UPR^mt^ in the *sod-2; sod-3* mutant (Fig. 4B). This result suggests that ICL-1 may counteract mitochondrial superoxide stress via a feed forward loop to increase UPR^mt^.

**Figure 4.**
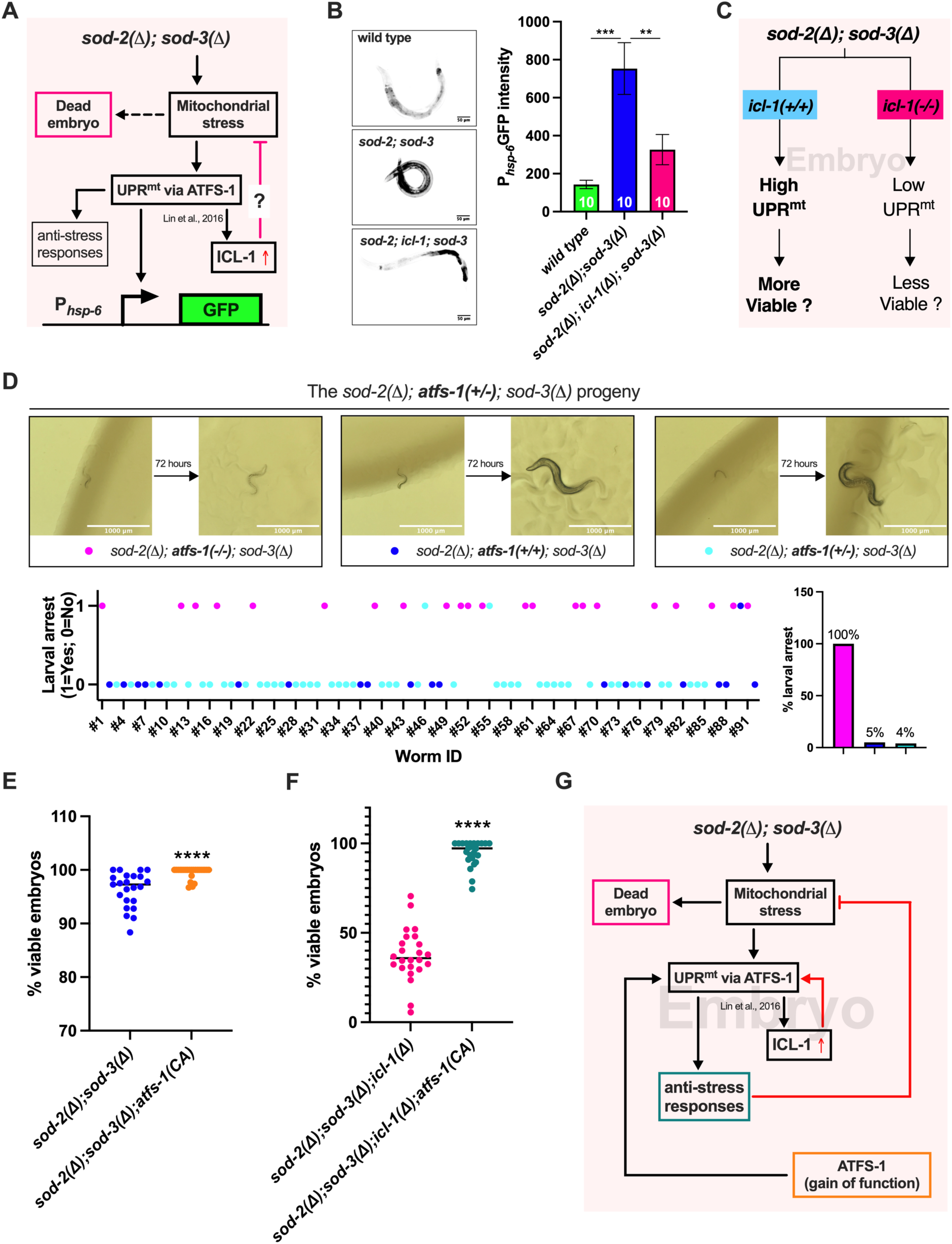
The interplay between the ICL-1 and UPR^mt^ in the absence of mitochondrial SODs engages ATFS-1. A. A diagram model proposing that ICL-1 induction will suppress the UPR^mt^ via ATFS-1. B. Representative pictures and Mean ± SEM of UPR^mt^ measured by the fluorescent intensity of P_*hsp-6*_GFP for wild type (mean = 143), *sod-2(ok1030); sod-3(tm760)* mutant (mean = 753) and *sod-2(ok1030); icl-1(ok531); sod-3(tm760)* mutant (Mean = 326) (n = 10 L4 stage hermaphrodites, *** *p*=0.0002 or ** *p*=0.0078 in one-way ANOVA). C. A diagram to propose that embryonic viability is correlated with UPR^mt^. D. Eliminating ATFS-1 in the *sod-2(ok1030) sod-3(tm760)* mutant causes larval arrest. A total of 92 progeny from multiple (n = 7) *sod-2(ok1030) atfs-1(+/gk3094) sod-3(tm760)* hermaphrodites are expected. There was no genotyping result for progeny #53, so the outcome of #53 is not annotated. The (-/-) vs. (+/+) vs. (+/-) count for *atfs-1* is 22 vs. 20 vs. 49. As for *atfs-1* in the *sod-2 sod-3* mutant, all 22 *atfs-1(-/-)* mutants (magenta dots) are larval arrest; 1 out of 20 *atfs-1(+/+)* wild type (blue dots) is larval arrest; and 2 out of 49 *atfs-1(+/-)* heterozygotes (cyan dots) are larval arrest. E. Mean ± SEM of percentage viable embryos from the *sod-2(ok1030) sod-3(tm760)* double null mutant and the *sod-2(ok1030) atfs-1gf(et18) sod-3(tm760)* (n = 24 animals from 3 independent trials, **** *p*<0.0001 in unpaired two-tail *t*-test). F. Mean ± SEM of percentage viable embryos from the *sod-2(ok1030) icl-1(ok531) sod-3(tm760)* triple null mutant and the *sod-2(ok1030) icl-1(ok531) atfs-1gf(et18) sod-3(tm760) atfs-1* (n = 24 worms from 3 independent trials, **** *p*<0.0001 in unpaired two-tail *t-*test). G. A diagram to summarize that (1) UPR^mt^ induces gene expression of *icl-1* as reported by Nargund et al. in 2015 (Nargund *et al*., 2015) and Lin et al. in 2016 (Lin *et al*., 2016); (2) ICL-1 induction may stimulate UPR^mt^ other than suppress it (see Discussion for details); (3) Other anti-stress pathways activated by UPR^mt^ may play a significant role in suppressing mitochondrial stress and subsequently supporting embryonic development in the absence of mitochondrial SODs.

### Activation of UPR^mt^ protects embryonic development in the absence of mitochondrial SODs

The UPR^mt^ can activate systematic protection against mitochondrial stresses (Lin *et al*., 2016; Nargund *et al*., 2012) and thus we investigated whether UPR^mt^ is protective for embryonic viability under superoxide stress associated with SOD elimination from mitochondria (Fig. 4C).

ATFS-1 is the key transcription factor that mediates the UPR^mt^. We crossed the *sod-2; sod-3* double null mutant with the *atfs-1* null mutant to examine the potential role of UPR^mt^ in counteracting mitochondrial superoxide stress. A subgroup of F2 generation that carries the *sod-2(-/-); **atfs-1(+/-);** sod-3(-/-)* genotype is viable and can produce the F3 generation, exhibiting a normal Mendelian inheritance ratio for the *atfs-1* mutant alleles: wild type vs. heterozygous vs. homozygous mutant = 20:49:22 (Fig. 4D). However, the *sod-2(-/-); atfs-1(-/-); sod-3(-/-)* triple null mutant F3 animals exhibited a 100% larval arrest phenotype, while the other two genotypes, *sod-2(-/-); atfs-1(+/-); sod-3(-/-)* or *sod-2(-/-); atfs-1(+/+); sod-3(-/-)* only have 4% or 5% larval arrest (Fig. 4D), respectively. The larval arrest phenotype suggests that *atfs-1* is essential to protect the healthy development of progeny under mitochondrial superoxide stress.

The F3 *sod-2; atfs-1; sod-3* triple null mutant animals only exhibited the larval arrest phenotype but did not exhibit significant embryonic lethality phenotype, which is most likely due to maternal effect rescue of residual ATFS-1 from the heterozygous parent. In the next set of experiments, we further assessed the role of *atfs-1* (or UPR^mt^) in counteracting mitochondrial superoxide stress by crossing the *atfs-1(et18)* constitutively active (CA) mutant (Rauthan et al., 2013) to either the *sod-2; sod-3* double null mutant or the *sod-2; icl-1; sod-3* triple null mutant. As shown by the earlier set of experiments, the *sod-2; sod-3* double null mutant has only slightly reduced embryonic viability (Fig. 2C); the *atfs-1(et18)* constitutively active mutation can significantly improve its embryonic viability from 96% to 99.5% (Fig. 4E). More strikingly, the *atfs-1(et18)* constitutively active mutation can improve the embryonic viability of the *sod-2; icl-1; sod-3* triple null mutant from 38% to 95% (Fig. 4F).

Together, ATFS-1 genetic data strongly suggest that the UPR^mt^ plays a prominent role in counteracting superoxide stress associated with the absence of mitochondrial SOD. The induction of *icl-1* expression may enhance the UPR^mt^ to counteract mitochondrial superoxide stress (Fig. 4G).

## DISCUSSION

Here we report that the *C. elegans* glyoxylate shunt enzyme ICL-1 is required for the survival of the embryo under mitochondrial superoxide stress. The glyoxylate shunt in *C. elegans* requires two key enzymes that are encoded by a single gene named *icl-1*. Knocking out Eliminating all mitochondrial SOD, but not eliminating cytosolic SOD, increases *icl-1* gene expression and, importantly, the *icl-1* gene induction is required for the survival of the mitochondrial *sod* null mutants. Consistent with the mitochondrial *sod* mutant findings, several mitochondrial ETC mutants, which are correlated with mitochondrial oxidative stress (van der Bliek *et al*., 2017), also have significant drop of brood size in the absence of ICL-1 (Fig. s1).

ICL-1 is known to protect against more than mitochondrial superoxide stress. Metabolic rewiring by the glyoxylate shunt can also promote synthesis of trehalose to increase the resistance to desiccation (Erkut et al., 2016). In our studies we cultured animals under standard nematode growth conditions, so the protective role of the glyoxylate shunt against mitochondrial superoxide stress is less likely due to the regulation of trehalose production.

ATFS-1, the core component of UPR^mt^, is a transcription factor mediating *icl-1* gene expression (Lin *et al*., 2016; Nargund *et al*., 2015), which indicates the involvement of ICL-1 in stress defenses. However, *icl-1* gene expression is also regulated by metabolic signaling pathways, including the upregulation of *icl-1* gene expression by starvation (Baugh et al., 2009) or disrupting the insulin signaling pathway (Depuydt et al., 2014; Hibshman et al., 2017; Murphy et al., 2003; Penkov et al., 2020; Senchuk *et al*., 2018) (Murphy, C. T. et al. 2003 Nature 424, 277–283; Shen, E. Z. et al. 2014 Nature 508, 128–132); and the requirement of lipid metabolism related NHR-49 transcription factor for *icl-1* expression (Dasgupta et al., 2020; Goh et al., 2018; Van Gilst et al., 2005; Zuryn *et al*., 2010). The biology of differential regulation of *icl-1* gene expression is quite poorly understood, but clearly the enzyme centrally interacts with cell metabolic defenses and energy metabolism.

Consistent with the previous report that *sod-2* null mutation by itself activates UPR^mt^ (Wu *et al*., 2018), our result demonstrates that *sod-2; sod-3* double null mutations can also activate UPR^mt^. The activation of UPR^mt^ could be due to the damage of mitochondrial function by superoxide, which is reflected by the lower mitochondrial inner membrane potential (Δ**Ψ**_m_) in *sod-2; sod-3* double null mutant as compared to the wild type (Fig. 3A-B).

Interestingly, the *icl-1* null mutation significantly reduces the activation of UPR^mt^ by mitochondrial superoxide stress, even though the glyoxylate shunt can theoretically reduce the NADH input to ETC by 6-fold to reduce superoxide production. A reasonable explanation is that the absence of glyoxylate shunt can increase the Δ**Ψ**_m_, a key driving force to prevent UPR^mt^ activation by reducing ATFS-1 migration into the nucleus (Melber and Haynes, 2018; Nargund *et al*., 2012; Rolland et al., 2019). However, our TMRE-based measurements fail to detect any significant increase/decrease of Δ**Ψ**_m_ upon *icl-1* knockout. Although the two potential effects of eliminating ICL-1, increased NADH input to ETC and increased superoxide production from ETC, might counteract to each other resulting in the no change of Δ**Ψ**_m_.

The embryonic viability results of *atfs-1(CA)* mutant further support the essential role of the UPR^mt^ in counteracting mitochondrial superoxide stress. Worms without both ICL-1 and mitochondrial SOD only have about 40% embryonic viability, however, a gain-of-function mutated ATFS-1 can increase the viability up to average 95%. These data indicate the protective mechanism is likely not directly executed by ICL-1 but most likely by genes that are upregulated by ATFS-1 to increase mitochondrial quantity/quality, improve resistance to oxidants or remodel metabolism (Lin *et al*., 2016). Overall, our exploration of the molecular mechanism by which ICL-1 counteracts mitochondrial superoxide stress reveals that ICL-1 is part of a feed forward loop for UPR^mt^.

It has been a puzzle for decades that why eliminating mitochondrial SODs causes neonatal lethality in mouse (Lebovitz *et al*., 1996; Li *et al*., 1995) or death in early adult age in flies (Duttaroy *et al*., 2003), but not in *C. elegans* (Van Raamsdonk and Hekimi, 2009; 2012). Since the glyoxylate shunt is not functional in mouse and fly, we speculate that restoring glyoxylate shunt in mouse or fly via expressing ICL-1 may prevent the neonatal lethality of mouse or fly in the absence of mitochondrial SOD since ICL-1 promotes UPR^mt^ activation and UPR^mt^ activation confers protection against mitochondrial superoxide stress to allow the survival of *C. elegans*. Differences between *C. elegans* UPR^mt^ vs mouse and fly UPR^mt^ might merit further investigation to extend understanding.

Our data suggest that ICL-1 plays a positive role in stimulating the UPR^mt^ mediated by ATFS-1. ATFS-1 is vital for embryonic development in the absence of mitochondrial SOD, and moreover, the constitutive active ATFS-1 mutation can dramatically improve embryonic viability of the *C. elegans* mitochondrial *sod* null mutants even in the absence of ICL-1. Wu et al. reported that eliminating ATFS-1 can cause significant larval arrest to mitochondrial ETC mutants, *clk-1* and *isp-1* (Wu *et al*., 2018). Meanwhile, both *clk-1* and *isp-1* mutants can cause significant upregulation of *icl-1* gene expression (Senchuk *et al*., 2018). These discoveries along with this study indicate that ICL-1 may play an important role in assisting UPR^mt^ to protect the programed embryonic development against mitochondrial stress.

The strong connection of ICL-1 to UPR^mt^ in counteracting mitochondrial superoxide stress suggests that ICL-1 can be used as a synthetic tool in treating diseases that are related to mitochondrial superoxide stress. For example, non-inheritable viral delivery of *icl-1* mRNA can potentially treat diseases caused by mutations on ETC complex I. ICL-1 can reduce the NADH-fueled superoxide production by the ETC complex I and promote the electron transport from ETC complex II. Since ETC complex II is less likely producing superoxide (Brand, 2010), the ICL-1 therapy might restore energy production with much lower risk of mitochondrial superoxide stress.

## METHODS AND MATERIALS

### Strains and maintenances

All strains are maintained on NGM (nematode growth medium) plate seeded with OP50-1 *E. coli* in a 20°C incubator. All *sod* mutants and the *icl-1* mutant have been outcrossed to the N2 wild type (PD1073 subclone) strain for at least 4 times. Strains used in this study include N2, CW152: *gas-1(fc21)*, MQ1333: *nuo-6(qm200)*, MQ887: *isp-1(qm150)*, ZB4570: *sod-2(ok1030); sod-3(tm760)*, ZB4572: *sod-1(tm783) sod-5(tm1146)*, ZB4665: *sod-2(ok1030); sod-1(tm783) sod-5(tm1146); sod-3(tm760)*, ZB4668: *icl-1(ok531)*, ZB4675: *sod-2(ok1030); icl-1(ok531); sod-3(tm760)*, SJ4100: *zcIs13[hsp-6::GFP]*, ZB4777: *sod-2(ok1030); sod-3(tm760); zcIs13[hsp-6::GFP]*, ZB5114: *sod-2(ok1030); icl-1(ok531); sod-3(tm760); zcIs13[hsp-6::GFP]*, ZB4680: *atfs-1(gk3094)*, QC118: *atfs-1(et18)*, ZB5018: *sod-2(ok1030); atfs-1(et18); sod-3(tm760)*, and ZB5113: *sod-2(ok1030); atfs-1(et18) icl-1(ok531); sod-3(tm760)*.

### RT-qPCR

All worms are harvested into TRIzol (Invitrogen) at the L4 stage and quickly frozen in liquid nitrogen. We performed RNA extraction and RT-qPCR as previously described (Laranjeiro et al., 2017; Laranjeiro et al., 2019). Briefly, after freeze-thaw cycles with liquid nitrogen/37 °C heat block, we extracted total RNA following the manufacturer’s instructions (Invitrogen) and synthesized cDNA using the SuperScript III First-Strand Synthesis System (Invitrogen). We performed quantitative PCR using diluted cDNA, PerfeCTa SYBR Green FastMix (Quantabio) and 0.5 μM of gene-specific primers in a 7500 Fast Real-Time PCR System (Applied Biosystems), calculating relative expression using the ΔΔCt method (Livak and Schmittgen, 2001) with *cdc-42* and Y45F10D.4 as reference genes (Hoogewijs et al., 2008). We used the following primer pairs: *icl-1*, ACGTTGCTGGAGACGAGAAG/AGAAGTCTCAAACGGCGAGG (amplification efficiency 0.97); *cdc-42*, CTGCTGGACAGGAAGATTACG/CTCGGACATTCTCGAATGAAG (amplification efficiency 1.03); Y45F10D.4, GTCGCTTCAAATCAGTTCAGC/GTTCTTGTCAAGTGATCCGACA (amplification efficiency 0.94).

### Brood size and embryonic viability assay

All worms are singled out into individual plate at the L4 stage and transferred to a new plate in each day until no more eggs or progeny appearing on the plate. The hatched progeny and dead embryos are counted in the old plate at least 24 hours (48 hours @ 20 °C for the mitochondrial *sod* mutants) after transferring the mother worm to a new plate.

### Membrane potential measurement

L4 stage hermaphrodites was placed into 35 mm NGM agar plates containing 100 μM TMREE (ENZO, Cat. #: ENZ-52309) in the bacteria mix. After 24 hours feeding, worms were immediately imaged under the confocal microscope.

### Paraquat toxicity

The age synchronized (by picking the L4 stage hermaphrodites) worms were placed into 60 mm NGM agar plates containing 2 mM paraquat (Sigma, Cat. #: 856177) in adult day 1. The viability of worms was recorded in the following days.

### UPR^mt^

The UPR^mt^ is carried out with quantification of GFP fluorescence expressed under the promoter of *hsp-6*. Images are taken on a spinning disc confocal microscope. The mean intensity of GFP among the whole body of L4 stage hermaphrodites is quantified with ImageJ. The representative pictures are color inverted in ImageJ.

### The DIC picutres

DIC pictures for eggs collected from plates are taken on a Zeiss compound microscope.

### The icl-1 null or atfs-1 null genotyping

A three primers strategy is used for genotyping. A forward primer (FWD) binds to one side out of the deletion region, a reverse primer (REV) binds to the other side out of the deletion region, and the 3^rd^ primer binds to the deletion region. The sequence of these primers are listed as below: *icl-1* FWD 5’-GTTGAAGGATGGAGAGCAGTTGTAG-3’; *icl-1* REV 5’-CAAACGTGACTATACCGTCGAAGATG-3’; *icl-1* 3^rd^ 5’-TCTGTTGATTCTCTTCGCCAATTCC-3’; *atfs-1* FWD 5’-CAGTGAGGCATAGAACGTGAGAATTG-3’; *atfs-1* REV 5’-GGAAAGTGGGTGGTGTGATGTTATT-3’; *atfs-1* 3^rd^ 5’-GTTGCTGATGACCATGTGGATGT-3’.

### Larval arrest pictures

Each L1 progeny is placed in an individual plate and the first picture is taken with a Revolve microscope in about 18 hours after isolation, then the second picture is taken 72 hours after the first picture. Images are processed by ImageJ for figure plotting.

### Figure plotting and statistics

All figures are plotted with GraphPad. The statistics tools in GraphPad are used to analyze the significance of differences. No power analysis is used for all experiments, but multiple repeats have been performed for all experiments.

## ACKNOWLEDGEMENTS

We would like to thank Dr. Rachel M. Woodhouse from University of Sydney for providing comments to the manuscript. Some strains were provided by the CGC, which is funded by NIH Office of Research Infrastructure Programs (P40 OD010440). This study is supported by NJCCR Postdoc Fellowship Award (#DHFS16PPC070) to G.W.

## AUTHOR CONTRIBUTIONS

G.W., S.L. and R.L. performed experiment, analyzed the data, and plotted the figures. G.W., J.V-R and M.D. designed the experiment and wrote the manuscript.

## DECLARATION OF INTERESTS

The authors declare no competing of interests.

## Notes

### Competing Interest Statement

The authors have declared no competing interest.

### Summary of Updates

Figure 4 revised; author email updated.

