## Supplementary material for "The glyoxylate shunt protein ICL-1 protects from mitochondrial superoxide stress through activation of the mitochondrial unfolded protein response": Figure s1

**Supportive figures:**

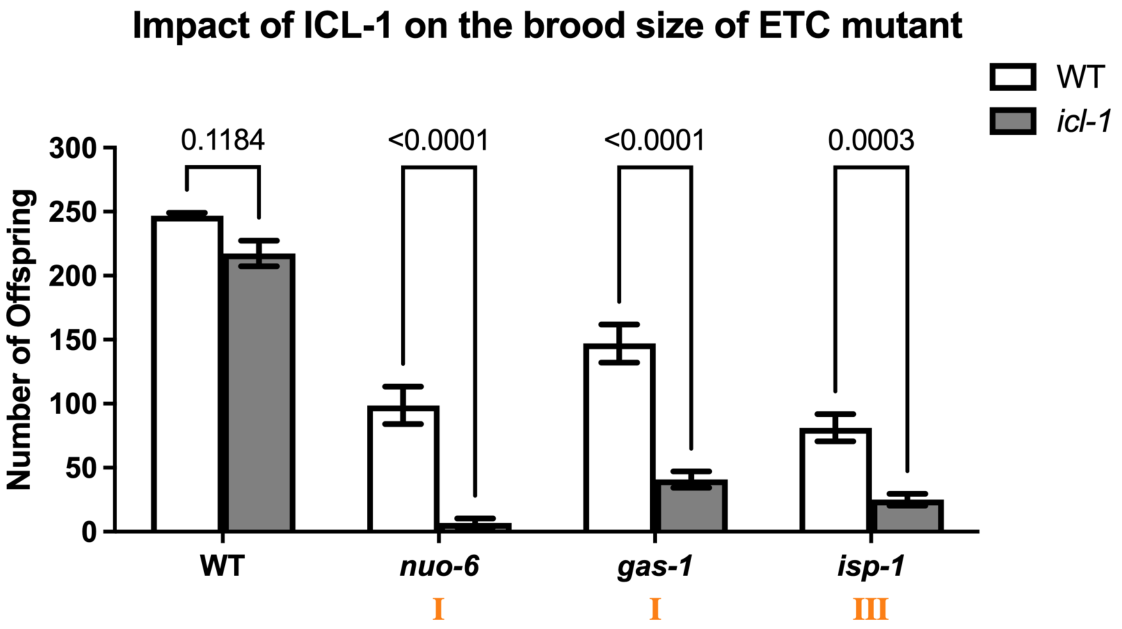


**Figure s1. Eliminating ICL-1 further reduces the brood size of some ECT mutants that are correlated with oxidative stress.**

Brood size of mitochondrial ETC mutant with/without *icl-1* null mutation. n = 15 hermaphrodite parents from three independent trials. The numbers on top of each paired columns are *p* value calculated from two-way ANOVA with Šídák's multiple comparisons test. The annotated Roman numerals (I and III) indicate mitochondrial electron transport chain (ETC) complexes.
